# Tactile suppression stems from sensation-specific sensorimotor predictions

**DOI:** 10.1101/2021.07.04.451060

**Authors:** Elena Führer, Dimitris Voudouris, Alexandra Lezkan, Knut Drewing, Katja Fiehler

## Abstract

The ability to sample sensory information with our hands is crucial for smooth and efficient interactions with the world. Despite this important role of touch, tactile sensations on a moving hand are perceived weaker than when presented on the same but stationary hand.^1–3^ This phenomenon of tactile suppression has been explained by predictive mechanisms, such as forward models, that estimate future sensory states of the body on the basis of the motor command and suppress the associated predicted sensory feedback.^4^ The origins of tactile suppression have sparked a lot of debate, with contemporary accounts claiming that suppression is independent of predictive mechanisms and is instead akin to unspecific gating.^5^ Here, we target this debate and provide evidence for sensation-specific tactile suppression due to sensorimotor predictions. Participants stroked with their finger over textured surfaces that caused predictable vibrotactile feedback signals on that finger. Shortly before touching the texture, we applied external vibrotactile probes on the moving finger that either matched or mismatched the frequency generated by the stroking movement. We found stronger suppression of the probes that matched the predicted sensory feedback. These results show that tactile suppression is not limited to unspecific gating but is specifically tuned to the predicted sensory states of a movement.

## Results

We investigated whether tactile suppression stems from sensorimotor predictions that estimate future sensory states of a movement or whether it can be better explained by unspecific gating that leads to a blanket reduction in tactile sensitivity, and thus does not distinguish between predicted and unpredicted sensory feedback. Tactile suppression is commonly tested by applying externally-generated tactile probes on a limb that is actively moving, a well-established paradigm that allows quantifying tactile suppression at various body parts and timepoints.^1,3,6^ Although this phenomenon has been attributed to predictive mechanisms, such as internal forward models,^4^ other accounts claim that it may stem from an unspecific gating that does not distinguish between predictable and unpredictable sensory states, but instead cancels out any tactile probes arising on the moving limb.^5,7^ Despite evidence that a movement-induced reduction in tactile sensitivity is accompanied by a downregulation of neural activity in the primary somatosensory and motor cortices already *before* movement onset,^8^ whether and how this phenomenon is related to prediction and expresses itself on the perceptual level is still obscure.

Here, we aimed to investigate the role of sensorimotor predictions in movement-induced changes in tactile sensitivity. To determine sensation-specific suppression effects, we manipulated the congruency between the predicted sensory feedback and the external vibrotactile probes. Participants performed single stroking movements across two textured objects at a designated speed. Depending on the spatial frequency of the object’s texture, they experienced either a low or a high frequency vibration on their fingertip. The two frequencies were chosen in a way that different mechanoreceptors were stimulated.^9,10^ The objects were presented in a blocked manner, so that the vibrating feedback from the movement was fully predictable. Vibrotactile probe stimuli were applied to the stroking finger around movement onset; their frequencies either matched (congruent condition) or mismatched (incongruent condition) the frequency elicited on the finger during movement across the texture. This resulted in four conditions: object low/probe low, object low/probe high, object high/probe high, object high/probe low.

We calculated the difference in tactile detection thresholds (threshold_diff_) and tactile precision (precision_diff_) for each participant between each stroking condition and a respective baseline (detection of low-frequency and high-frequency probe at rest, see Figures 1A and 1B). Positive values of threshold_diff_ and precision_diff_ indicate stronger tactile suppression and worse detection precision during movement, respectively. If tactile suppression stems from a predictive mechanism, we expect stronger suppression (increased threshold_diff_) in congruent than in incongruent conditions, as the probe frequencies would match the movement-related predictions. If suppression is also associated with a reduction in detection precision,^11^ detection responses should be less precise in congruent than incongruent conditions. If, on the other hand, tactile suppression stems from a general gating mechanism, then we should observe similar suppression across all conditions (comparable threshold_diff_).^5,7^

**Figure 1.**
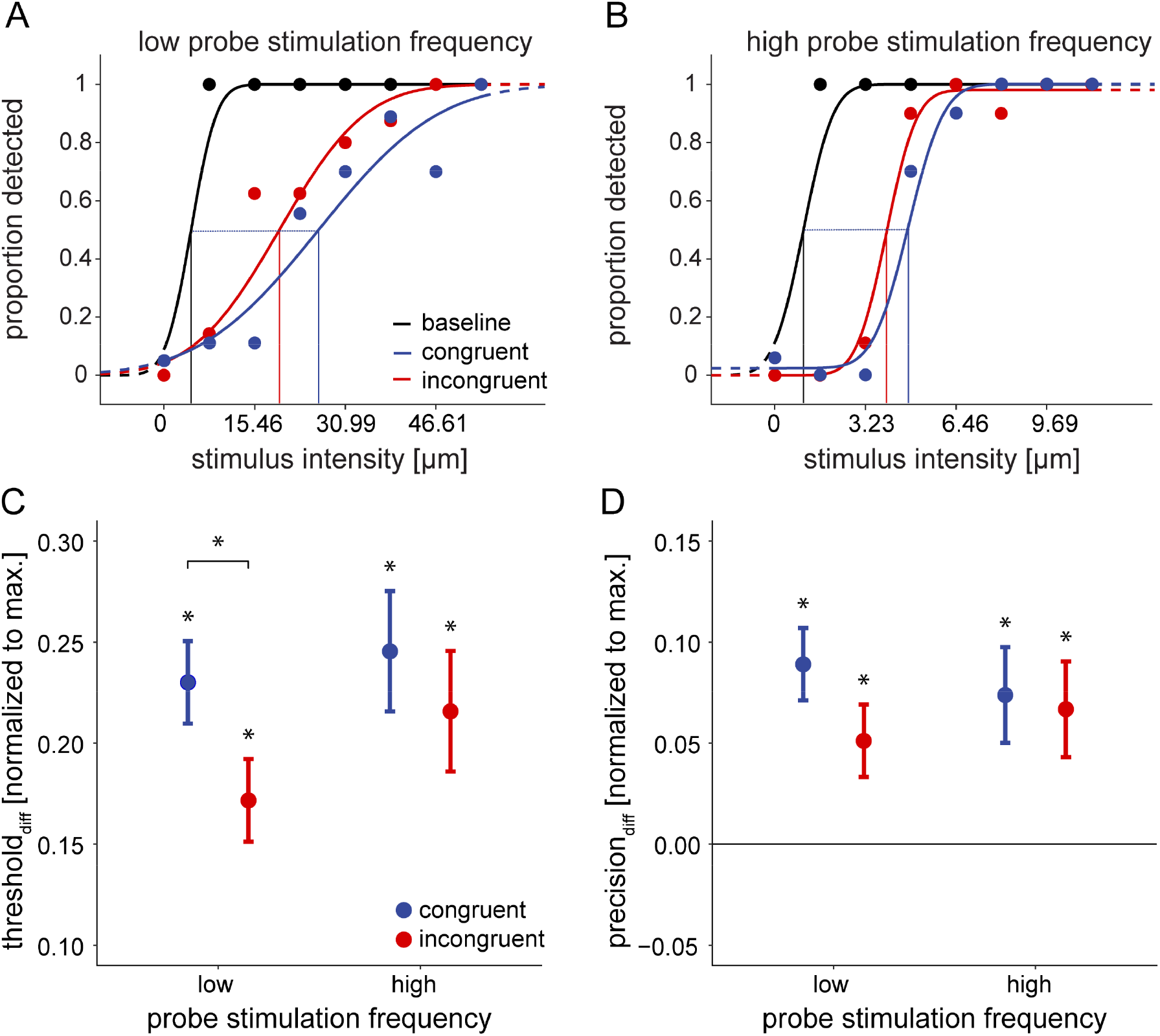
Tactile Suppression in the Movement Conditions. *Note*. Example psychometric functions of a single participant for low (A) and high (B) frequency probes. Each panel shows psychometric functions for the baseline and the two movement conditions. The difference between the baseline and the movement conditions is exemplified by the dotted line representing threshold_diff_ for the congruent movement condition. Differences in detection of the probe stimulation between the baseline and the movement conditions as measured by the change in detection thresholds (C) and detection precision (D), averaged across all participants. All values are normalized to the maximum possible suppression at the respective probe frequency. Greater values indicate impaired sensitivity to probes, i.e., tactile suppression, in the movement conditions, while zero indicates no difference from the baseline. C. Tactile suppression took place in all movement conditions and was generally greater in congruent compared to incongruent conditions. D. Detection precision was hampered in all movement conditions. There were no differences in the detection precision of congruent and incongruent probe stimulations. ^a^The error bars display 95% Cousineau-Morey confidence intervals.^14,15^

First, to determine whether tactile perception was affected during the movement as compared to rest, we performed t-tests against zero on threshold_diff_ and precision_diff_. In line with previous findings,^12,13^ tactile stimuli were suppressed in all movement conditions, as evidenced by detection thresholds_diff_ that were significantly greater than zero for both low frequency congruent, *t* (31) = 6.48, *p* < .001, *d* = 1.45, and incongruent, *t* (31) = 6.27, *p* < .001, *d* = 1.11, conditions, as well as high frequency congruent, *t* (31) = 7.63, *p* < .001, *d* = 1.35, and incongruent, *t* (31) = 7.30, *p* < .001, *d* = 1.29, conditions (see Figure 1C). Detection precision was hampered during movement compared to rest in all conditions, for low frequency congruent, *t* (31) = 4.56, *p* < .001, *d* = .81, and incongruent, *t* (31) = 3.12, *p* = .004, *d* = .55, conditions, as well as for high frequency congruent, *t* (31) = 3.45, *p* = .002, *d* = .61, and incongruent, *t* (31) = 3.77, *p* < .001, *d* = .67, conditions (see Figure 1D).

Second, we examined whether suppression was stronger in congruent than incongruent conditions, which would provide strong evidence for the involvement of predictive mechanisms. To this end, we compared threshold_diff_ and precision_diff_ using 2 (object: low vs. high) by 2 (probe: low vs. high) univariate, repeated-measures ANOVAs. Neither the type of the object, *F* (1,31) = 0.64, *p* = .43, η_G_^2^ = .002, nor the frequency of the probe, *F* (1,31) = 0.75, *p* = .39, η_G_^2^ = .007, influenced the thresholds_diff_. Importantly, tactile suppression was stronger for congruent compared to incongruent conditions as evidenced by a significant interaction, *F* (1,31) = 6.31, *p* = .017, η_G_^2^ = .016. This difference was particularly systematic for thresholds_diff_ of low frequency probes, *t* (31) = 2.92, *p* = .003, *d* = .52, and less so for high frequency probes, *t* (31) = 1.01, *p* = .16, *d* = .180 (see Figure 1C). No systematic differences were found in the precision_diff_, with neither a main effect of the object type, *F* (1,31) = 1.01, *p* = .32, η_G_^2^ = .005, nor of the probe frequency, *F* (1,31) < 0.01, *p* = .99, η_G_^2^ < .001, nor an interaction of the two, *F* (1,31) = 2.67, *p* = .11, η_G_^2^ = .011 (see Figure 1D).

Last, to test if the differences in tactile suppression found between the movement conditions may be due to differences in the movement itself, rather than to a prediction of the sensory states of the movement, we compared several kinematic parameters using 2 (object: low vs. high) by 2 (probe: low vs. high) univariate, repeated-measures ANOVAs. No systematic differences were found for the reaction time, all *F* (1,31) ≤ 3.34, *p* ≥ .08, η_G_^2^ ≤ .002, the movement duration, all *F* (1,31) ≤ 1.13, *p* ≥ .3, η_G_^2^ ≤ .005, the average force exerted during the movement, all *F* (1,31) ≤ 3.00, *p* ≥ .09, η_G_^2^ ≤ .010, the average movement velocity, all *F* (1,31) ≤ 1.57, *p* ≥ .22, η_G_^2^ ≤ .007, as well as the stimulation time, all *F* (1,31) ≤ 2.12, *p* ≥ .16, η_G_^2^ ≤ .007 (see Figure 2).

**Figure 2.**
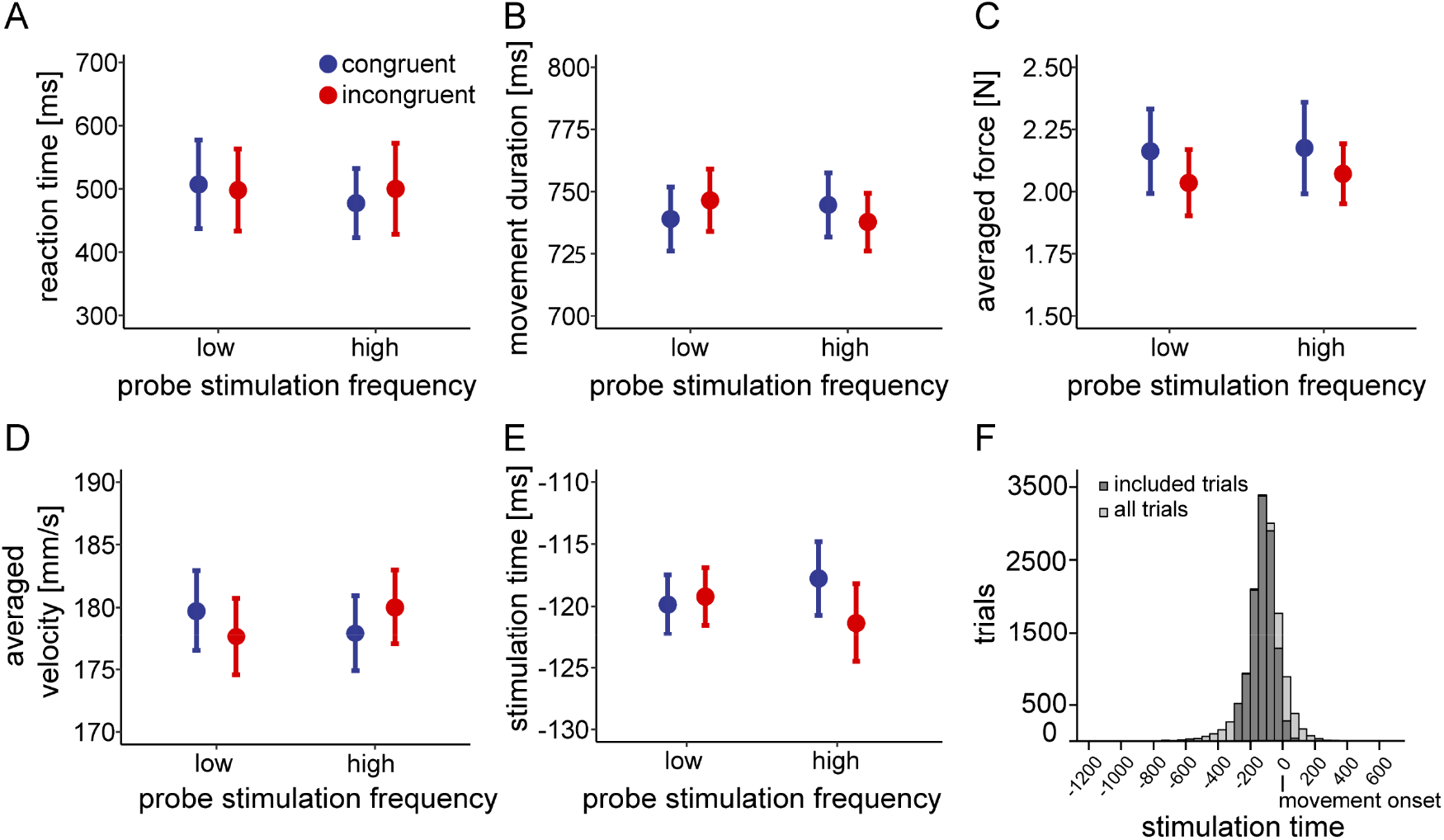
Kinematic Parameters in the Movement Conditions and Timing of the Probe Stimulation. *Note*. Kinematic parameters averaged across participants and movement conditions. A. Reaction time. B. Movement duration. C. Force averaged across the movement. D. Velocity averaged across the movement. E. Stimulation time. F. Stimulation times in relation to movement onset (0 ms). The stimulation times included in the analysis occurred no earlier than 300 ms before movement onset. To avoid backward masking of the probe by the texture, stimulations ending less than 150 ms before the finger reached the texture were excluded.^16^ None of these movement parameters differed between the conditions. ^a^The error bars display 95% Cousineau-Morey confidence intervals.^14,15^

In summary, congruent vibrotactile probes matching the predicted sensations caused by the movement across textured surfaces were subject to stronger suppression than incongruent probes. This suppression was reflected in the detection thresholds, but not in the detection precision. No significant differences of movement kinematics were found across conditions, excluding the possibility of systematic movement effects on the detection performance.

## Discussion

Suppression of externally-generated sensations shortly before and during voluntary movement is a well-established phenomenon.^17–19^ Suppressing sensations that are not caused by one’s own actions appears paradoxical because survival and adaptation require to enhance, and not to suppress, novel and potentially harmful sensations. Consequently, the mechanisms governing sensory suppression of external stimulations are still a matter of debate. On the tactile domain, it is claimed that the reduced sensitivity on a moving limb is caused by general cancellation policies,^5^ such as unspecific gating of reafferent signals. However, there is accumulating evidence that central, predictive mechanisms regulate suppression of tactile,^12,18^ auditory,^20–22^ and visual stimuli.^23,24^ Though, the neural and perceptual suppression effects do not necessarily reflect the same mechanism.^25^ We here set out to address this debate and show that tactile suppression of externally-generated sensations is modulated by predictive mechanisms.

To resolve whether suppression of external vibrotactile probes stems from specific sensorimotor predictions or an unspecific gating mechanism, we asked participants to stroke with their finger along two different textures, with the movement across each texture causing a distinct vibration on the stroking finger. Meanwhile, we presented probes of congruent or incongruent vibration frequencies on that finger. These probes served as a proxy for the sensory states associated with the movement.^12,18^ Consistent with previous work,^3,6,12,16^ we observed increased detection thresholds and hampered detection precision of tactile sensations around movement onset compared to detection at rest. Critically, congruent conditions led to stronger suppression of detection thresholds, but not of precision. This finding indicates that tactile suppression originates from a specific sensorimotor prediction about the future sensory states generated by the movement.

Sensorimotor control is thought to depend on the optimal integration of a sensorimotor estimate about the future sensory states of the system and of peripheral sensory input.^26^ Thus, when the state of the *environment* is uncertain, the system needs to harvest sensory input, and thereby feedback gains are upregulated while predictive control is compromised.^27^ However, when the state of *one’s own sensory system* is highly uncertain, sensory feedback is downweighed and reliance on sensorimotor estimates is greater.^28^ Accordingly, in the presence of sensorimotor predictions, sensory feedback, including any external probes, is downweighed to different extents, depending on the reliability of the prediction. This is reflected in attenuated responses of sensorimotor cortices to tactile probes on a limb shortly before its movement^8^ and during movement.^29^ Tactile suppression of external probe stimuli is also modulated by feedback demands, such that it is weaker when task-relevant feedback gains importance,^12,19,30^ and stronger when movement feedback becomes more predictable.^16^ This suggests that suppression depends on an interplay between predictive and feedback signals.^6^ However, no study hitherto has distinguished predictive from general suppression mechanisms, leaving the phenomenon open to different interpretations. Our findings can reconcile computational approaches of sensorimotor control with tuning of sensory processing, and thus contribute to resolving a debate about the origins of tactile suppression.

Our results are incompatible with the idea that suppression of external probe stimuli is due to an unspecific gating mechanism that is caused by peripheral reafferences, independent of sensorimotor predictions.^5,7^ Although a general gating mechanism could explain the observed suppression in all four movement conditions compared to rest, it cannot account for differences between congruent and incongruent conditions. This congruency effect cannot be attributed to differences in the movement either,^16,31,32^ as kinematic behaviour was similar between the conditions. With the present results we cannot determine the extent to which gating may contribute to the observed suppression effects. As tactile suppression does not depend on the strength of reafferent input^33^ and, importantly, can fall off completely during movement,^6^ even at moments when transmission of muscle spindle afferences is increased,^34^ some additional modulatory mechanism seems to be involved. Our present findings support the idea that tactile suppression stems from a specific prediction about the future sensory states of the movement, providing novel evidence for a predictive mechanism involved in this phenomenon. Thus, unspecific gating may not be a competing explanation for the copious accounts of action-related suppression of externally-generated stimulations,^6,12,19,30^ but rather a separate mechanism operating in addition to predictive suppression.

Suppression may also arise due to a reduction in the precision (reliability) of afferent information on the moving limb,^7^ as proposed in the active inference account.^11,35,36^ According to this account, a movement is generated to fulfil the sensory predictions expected from a specific action. Reduced precision of somatosensory feedback during the movement allows expressing sensory predictions that trigger the movement. This means that a reduction in the precision is a pre-condition to generate sensory predictions reflected in increased thresholds, i.e. stronger suppression. Here, we found that detection precision was indeed worse during movement compared to rest (baseline), in line with the active inference account. However, we did not find an effect of congruency. Descriptively, precision was worse in congruent than incongruent conditions, but this effect was not systematic. Therefore, the elevated detection thresholds in the congruent conditions are unlikely to be caused by poorer detection precision.^7,11^

Tactile suppression has also been explained through the opposing process theory,^5^ according to which predicted sensations may first be enhanced and later in time cancelled. This theory focuses on the perception of self-generated tactile events, and thus leaves open whether and how it applies to external probe stimuli serving as a proxy to measure predictive tactile suppression on that limb.^8^ While the opposing process theory draws evidence from the visual modality,^37^ it is at odds with work from the tactile domain showing predictive suppression of self-^38^ and externally-generated tactile sensations^3,19^ occurring up to 300ms before movement onset, i.e. lacking preceding enhancement.

Stimulating just before movement onset, and well before touching the textured surface, is a common method to measure tactile suppression during movement planning while controlling for influences of backward masking^2,16,39^ by the texture. Naturally, the perception of the probe may have been affected by moving across the initial, smooth part of the object, between the go position and the texture. This, however, cannot account for the differences between the congruent and incongruent conditions. It is therefore unlikely that the reported congruency effects are caused by the movements across the texture. Because the minimum interval between the onset of the probe and the moment when the texture was first touched (250 ms) far exceeded the threshold of temporal acuity in the tactile domain (50-70 ms),^40^ this early stimulation timing allowed for the perception of the probe stimulation and the texture as two distinct events.

The congruency effect was more pronounced for low than for high frequency probes. This could be due to the two types of mechanoreceptors that were targeted with the stimulation frequencies. With probes of 40 Hz we likely stimulated the Meissner corpuscles that have a receptive range of 10-100 Hz, and are most sensitive to vibrations of 40-60 Hz.^9,10^ The probes of 240 Hz lie in the response range of the Pacinian corpuscles that extends from 40-800 Hz,^9^ with their maximal sensitivity being between 200-300 Hz.^10^ It is conceivable that the higher specificity of the Meissner corpuscles associated with the low frequency probes led to a better match between the estimated and sensed vibration, and therefore to a more pronounced suppression effect.

We conclude that tactile suppression of externally generated stimuli on a moving limb is modulated in a sensation-specific manner, as it is stronger when the generated stimuli match the predicted sensory feedback. This speaks in favour of a predictive component involved in tactile suppression, in addition to tactile gating. Our findings do not only provide novel evidence for the origins and underlying mechanisms of this phenomenon; they also form the ground to unify the mechanisms underlying tactile suppression with those determining sensory tuning in other modalities, such as vision^23,24,41^ and audition.^42^ This contributes to a better understanding of the computational principles and neurobiological substrates of human sensorimotor control as well as of clinical phenomena related to predictive mechanisms, such as Parkinson’s disease,^43^ OCD,^44^ schizophrenia,^45,46^ and depression.^47^

## Methods

### Participants

A total of 48 participants completed the experiment and were compensated with either course credit or a payment of 8€/hour. Due to exclusion criteria listed below, 32 participants (23 women, 9 men, range 19-30 years, 22.56 +/-3.03) were included in the final sample (see Quantification and statistical analysis for the exclusion criteria and the sample size calculation). The experiment was approved by the local ethics committee at the Justus Liebig University Giessen and was in accordance with the Declaration of Helsinki from 2008. All participants provided their signed informed consent. Participants were right-handed according to the German translation of the Edinburgh Handedness Inventory^48^ (77.94 +/-21.48). Furthermore, participants reported no current neurological symptoms, no issues with stereopsis or colour vision and had normal or corrected to normal eyesight. Participants also did not report any injuries to the right index finger and had a two-point-discrimination threshold of at least 3 mm on the right index finger (2.34 +/-0.48 mm; Two-Point Discriminator, North Coast Medical, Inc., Morgan Hill, CA).

### Apparatus

The experiment was conducted using a 3D virtual environment, a force feedback device (PHANToM 1.5A, 3D Systems Inc., Rock Hill, SC) in connection with a force sensor (bending beam load cell LCB 130 and a measuring amplifier GSV-2AS, resolution .05 N, ME-Messsysteme GmbH), and a vibrotactile stimulation device (Engineering Acoustics, Inc., Casselberry, FL). Custom-made software (C++) was used to run the experiment, collect responses, and record the force and finger position every 3 ms. Participants sat in front of a table and connected their hand to the force feedback device using a custom-made plastic fingernail which was attached to the nail of their index finger with mouldable adhesive pads. This plastic fingernail was equipped with a concave magnetic element which fit snugly onto a matching metal ball point at the force feedback device. This setup allowed to keep the finger pad free for haptic exploration and permitted unrestricted movements within all six degrees of freedom within a workspace of 38 × 27 × 20 cm. Participants had no view of their finger connected to the force feedback device or the objects and instead viewed a spatially aligned 3D representation of the scene on a 22” monitor (120 Hz, 786 × 1024 pixel) by looking through a mirror using a pair of stereo glasses (3D Vision 2, Nvidia Corp., Santa Clara, CA). Visual stimuli associated with experimental control, such as questions and answers for the detection task, were presented on the monitor and corresponded to specific virtual positions in the workspace, so participants could navigate through the experiment and indicate answers by moving their finger, and thus the force feedback device, to those virtual buttons.

A vibrotactile stimulation device (tactor) was attached to the ventral part of the proximal phalange of the participants’ right index finger and was secured with medical tape. This tactor could present brief (100 ms) vibrotactile probes of eight varying intensities for each of the two probe frequencies that we used (40 Hz; 240 Hz). The peak-to-peak displacements of the stimuli ranged from zero (no stimulation) to 54.45 µm and to 11.31 µm (see Supplement) for probe frequencies of 40 Hz and 240 Hz, respectively, both in the baseline and movement blocks. We determined the stimulus intensities for the 40 Hz and 240 Hz probe frequencies in two pilot studies. In a first pilot study (*N* = 10) with identical intensities for both probe frequencies, we observed higher detection thresholds and higher suppression of low than high frequency probes. This resulted in a high number of threshold estimations that exceeded the range of presented stimulus intensities and negatively affected threshold estimations. Therefore, in a second pilot study (*N* = 6), we tested separate intensities for each probe frequency in order to obtain threshold estimations approximately halfway the range of the presented stimulus intensities for each probe frequency. White noise was played through over-ear-headphones to prevent participants from hearing the sound caused by the vibrotactile stimulation.

The textured objects were printed using a 3D PolyJet printer (Stratasys Ltd., Eden Prairie, MN; Objet30 Pro; printing resolution: 600 to 1600 dpi; material VeroClear). The objects were 209 mm long and 40 mm wide. The right half of the object had a texture with an even cube-wave pattern. The left half of the object was smooth to mitigate the risk of participants adapting to the texture over time. To create two distinct objects, a texture with a high spatial period of 5.08 mm, and another one with a smaller spatial period of 0.85 mm were chosen. When moving across these textures at a constant velocity of 0.203 m/s (average recorded speed 0.203 ± 0.032 m/s, range = 0.109 – 0.323 m/s), these spatial periods elicited vibrations with frequencies of 40 Hz and 240 Hz, respectively, on the participant’s fingertips (see Figure 3B).

**Figure 3.**
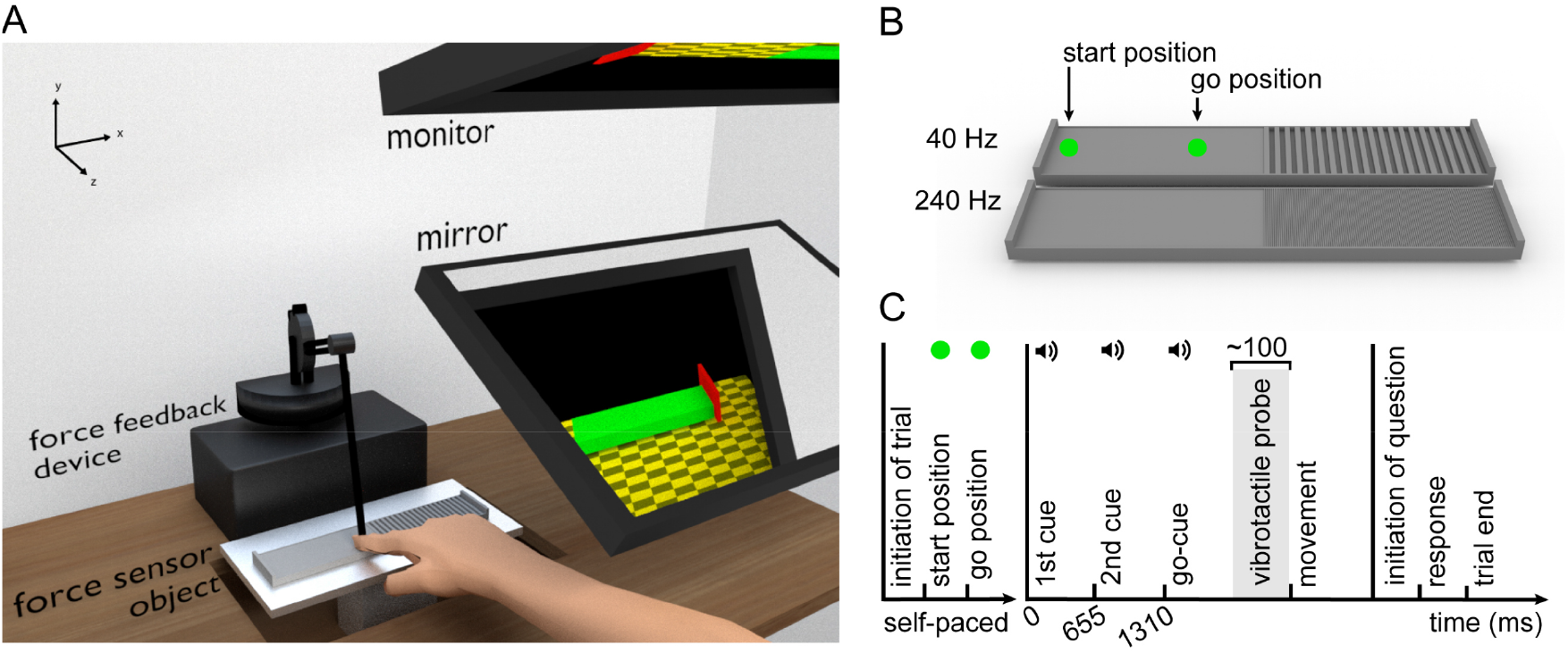
Experimental Setup, Textured Objects and Sequence of a Trial in the Movement Condition. *Note*. A. Illustration of the experimental setup. Participants were seated in front of a desk with the force sensor, the object and the force feedback device placed in front of them. A mirror blocked the view of the hand and the object. Participants viewed the scene presented to them on the monitor through the mirror. B. The low-frequency (40 Hz) and the high-frequency (240 Hz) objects. The start position at which participants initially made contact with the object and the go position from which participants began the stroking movements across the texture are marked with arrows. These two positions were represented to participants visually by green spheres on the monitor and were not discernible by touching the object. C. Sequence of a single trial during the movement conditions. Participants initiated each trial via a virtual button with the left hand. They were then presented with a green circle as a cue to move to the leftmost edge of the object (start position). Once they had reached this point, the green circle vanished and reappeared at the go position. As soon as participants reached the go position, the cue disappeared and a green, untextured cuboid representing the part of the object to the right of the start was shown. Simultaneously, three auditory cues spaced 655 ms apart were presented, the last of which was the go-cue. The vibrotactile probe (duration 100 ms) was triggered 100 ms before the median reaction time of the five previous trials. Once participants had finished their movement and reached the right end of the object, they could initiate the question regarding the detection of the vibrotactile probe via a central virtual button. After responding to this question, the trial was finished.

### Procedure

The experiment consisted of a sequence of four blocks, one for each movement condition (2 probe frequencies * 2 object frequencies), and of a single resting block that served as a baseline measure. To familiarize participants with the use of the force feedback device and the nature of the tactile stimulations, they first performed 20 mock trials of the baseline. This data was discarded and not included in the analysis. Next, half of the participants continued with the baseline whilst the other half completed the baseline at the very end of the experimental session. Before the first movement block, participants performed two blocks of practice trials to train their movement across a smooth object at the designated speed. Before the 2^nd^, 3^rd^, and 4^th^ movement block, they performed one practice block of eight trials. The order of the four movement conditions was pseudo-randomised with a latin square design resulting in four possible sequences, with each condition being equally likely to occur at each position.

The baseline test served to determine each individual’s tactile detection performance for each probe frequency. Participants placed their right hand comfortably on the table in front of them whilst the finger with the tactor rested on a pad to avoid the vibrations from reverberating through the table and distorting the measurement. In the baseline condition only, the participants’ left index finger was attached to the force feedback device to control the experiment and to limit possible effects on tactile perception from subsequent movements of the stimulated hand. Participants started each stimulation trial by bringing their left finger to a virtual button. After 100 ms, a probing tactile stimulus was delivered to their right index finger followed by a delay of 500 ms. Then, a question asking whether participants had felt a vibration was presented on the virtual monitor and could be answered with ‘yes’ or ‘no’ by selecting the respective virtual button with the left finger. During the baseline block, the right hand remained completely stationary.

The sequence of a single trial during the movement condition is depicted in Figure 3C. In each of the four movement conditions, the participants’ right index finger was attached to the force feedback device. They moved their finger across the textured objects at a constant speed of 0.203 m/s while detecting the vibrotactile probes. Upon initiating the trial by moving to a virtual button, a schematic representation of the object’s left and right outer border was presented, along with a green circle at the very left of the object (start position). As soon as participants moved their right finger to that circle, thus touching the real object in the workspace, the circle vanished and reappeared at the go position, 7 cm further to the right (∼3 cm before the textured area). Participants moved their finger at their own pace to that new position while keeping contact with the object. To prevent participants from drifting back again after having reached the go position, a force ‘wall’ was generated by the force feedback device to the left of this point. Once participants reached the go position, an untextured green block representing the remaining right part of the object appeared on the monitor followed by three auditory cues spaced 655 ms apart. This time interval of 655 ms corresponded to the time participants needed to travel the complete remaining distance of the object at an average speed of 0.203 m/s and thus aided the participants to adhere to this designated speed. Accordingly, participants started a smooth, continuous movement with the last of the three consecutive auditory cues (go-cue) and had to finish their movement at the time of an imagined fourth cue. Once they reached the end of the textured area, the object’s visual representation disappeared, and participants could initiate the detection question about the tactile stimulus via a central virtual button and respond as described in the baseline procedure, but now with their right hand.

To avoid any influence of the textured object and/or the movement itself on tactile suppression, we presented the probe stimuli shortly before movement onset, specifically 100 ms before the median reaction time of the preceding five movement trials.^29,31^ For the first five trials, for which no data was available to estimate the participants’ reaction time, the stimulation was presented 150 ms after the go-cue. For each probing frequency, we presented 70 trials with stimulation (each of the seven stimulation intensities repeated 10 times) and 20 trials with no stimulation in a randomized order. This resulted in a total of 90 trials for each block of the movement condition and 180 trials for the baseline that involved both probe frequencies.

Before the start of the first movement block, all participants performed two blocks of practice trials to train the stroking movements across an entirely smooth object at average speeds of 0.203 m/s. In the first of these two practice blocks, no probes were presented. Otherwise, the procedure corresponded to the trials of the movement condition but included visual feedback: During the movement, the finger position was represented by a vertical, red line. To give participants online feedback on their movement, vertical, blue lines moved across the scene at the required speed, allowing for variations in reaction time. Thus, if participants moved at a constant pace relative to these blue lines, they were adhering to the appropriate speed. The second practice block did not include the above-mentioned visual feedback and was similar to the movement conditions, except that we presented probes of an intermediate frequency (140 Hz) at varying intensities combined with a smooth object. This frequency of 140 Hz was used for two reasons: first, to avoid pre-emptively giving participants practice with one of the frequencies used during the movement blocks. Second, to avoid having to mix two frequencies within one practice block to be consistent with the upcoming movement blocks that contained probes of only one frequency each.

In the movement conditions and the practice blocks, trials were marked as invalid if participants moved either too fast (exceeding an average lateral velocity of 0.267 m/s) or too slow (falling below an average lateral velocity 0.140 m/s) along a section of the object (0.5 cm to the right of the go position until 1 cm to the left of the object’s right border). Trials were also marked as invalid when the go position was crossed before the go-cue was given. These invalid trials were repeated later by being randomly interleaved with the remaining trials of the block. On average, 9 +/-6% of trials were repeated in the four movement blocks. When an invalid trial occurred, as well as when the average stroking force fell below 1 N, the experimenter gave relevant verbal feedback: During the practice blocks, this feedback was given as needed. During the four movement blocks, the feedback was standardized and given after five mistakes had been made regarding the speed and the force, and after three mistakes when the movement onset was too early.

### Quantification and statistical analysis

The sample size was determined through an a priori power analysis based on the size of the congruency effect found in a previous pilot (*N* = 10). According to those data, to find an effect of congruency in the size of η_p_^2^ = .21, given α = .05 at a power (1-β) = .81, 31 participants were needed.^49^ The sample size was eventually set to 32 to allow for even randomization of the sequences of the four blocks.

The kinematic data obtained for each trial of the movement condition were processed using Matlab R2021a (The MathWorks, Inc., Natick, MA). Movement onset was defined by first determining the time when a position 0.5 cm to the right of the go position was crossed and then going backwards from this time to identify when the lateral velocity had first exceeded 0.67 mm/s. To determine the endpoint of the stroking movement, the Multiple Sources of Information method^50^ was applied to look into all frames when the finger applied force to the texture, using probabilistic criteria: a frame was more likely to be defined as the movement end the lower its velocity was and the closer its x-position was to the maximum (rightmost) x-position. The end of the movement was defined as the first frame in which this calculated endpoint was reached during the movement.

The average movement duration across trials in the four movement conditions as well as the kinematic parameters averaged across the timecourses are depicted in Figure 4. The reaction time refers to the time between the auditory go-cue and the movement onset and the movement duration to the time between the onset and the end of the movement. The force was calculated as follows. First, we filtered the force data to counteract noise: Any frames with force values exceeding 12 N or falling below –2 N were removed. To deal with oscillations, we applied a simple moving average filter with a window size of 60 ms. We then averaged the force data across the entire movement, i.e. from movement onset until movement end. The velocity was defined as the difference in x-position from one frame to the next and divided by the sampling interval of 3 ms. As the force, the velocity data was averaged across the entire movement.

**Figure 4.**
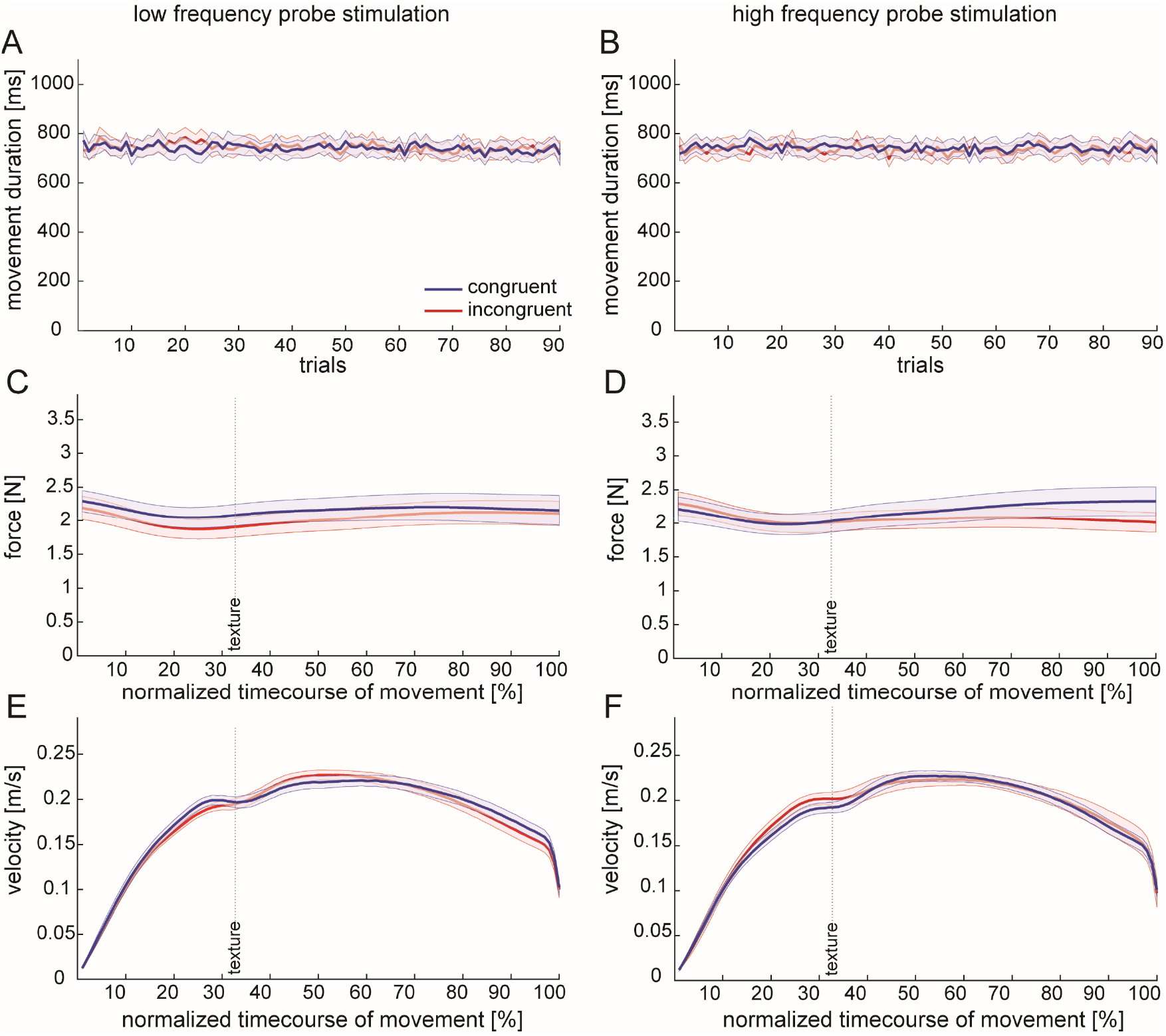
Movement Duration and Kinematic Parameters across Normalized Movement Timecourses. *Note*. The movement duration averaged across participants for each trial of the conditions involving (A) the low frequency probe stimulation and (B) the high frequency probe stimulation. The kinematic parameters averaged across participants for the normalized timecourse of the movement. C. Force (low frequency probe stimulation conditions). D. Force (high frequency probe stimulation conditions). E. Velocity (low frequency probe stimulation conditions). F. Velocity (high frequency probe stimulation conditions). ^a^The shaded error bars display 95% Cousineau-Morey confidence intervals.^14,15^

To avoid any effects of backward masking by the texture, only trials in which the stimulation ended at least 150 ms before participants reached the texture were included in the analysis of the tactile detection task.^2,16,39,51^ Likewise, only trials with stimulations starting no earlier than 300 ms before movement onset were included, to avoid probing suppression at a moment that is too early to occur.^3,19^ Based on these criteria, 84 +/- 10% of trials were included on average per participant. One participant was excluded because 50% of the trials had to be removed based on the two above-mentioned criteria. The average timespan between the onset of the stimulation and the moment when the finger reached the onset of the texture was 362.93 +/- 23.73 ms, while the probing stimulus was presented on average 113.87 +/- 12.39 ms before movement onset. Thus, our probing stimuli were presented during the time window when tactile suppression already occurs^3,19^ (see Figure 2F).

The responses to the tactile detection task were fitted as cumulative Gaussians, using the maximum likelihood procedure implemented in the psignifit 4 toolbox^52^ for Matlab. Separate functions were fitted for each of the four movement conditions and each participant. Two additional psychometric functions were fitted for responses to the low and the high frequency probes during the baseline. Due to a technical bug in the baseline condition, the first period of the sinusoidal probe stimulation for trial *n* was automatically set to the probe frequency of the trial *n-1*. Therefore, we fit baseline psychometric functions by using only the second of two consecutive trials that had the same frequency (50 +/- 3% and 50 +/- 4% for low and high probe stimulation frequency). For each of these functions, we calculated the detection threshold as the intensity that had a 50% probability of being detected, and the detection precision (slope) as the difference between the detection threshold and the intensity detected with a probability of 84%. To account for differences across individual baseline detection sensitivities, we subtracted each participant’s baseline detection threshold and precision from their respective values of each of their movement blocks, separately for low and high frequencies. This resulted in four detection_diff_ and four precision_diff_ values per participant, with higher values representing stronger suppression during movement compared to rest. To rule out any poorly fitted data, participants with detection thresholds exceeding the range of stimulation (*N* = 9) or with false alarm rates of 20% or more (*N* = 6) were excluded.

To examine whether suppression occurred during movement compared to rest, detection_diff_ and precision_diff_ were tested against zero using two-tailed t-tests. Possible effects of congruency on tactile suppression were examined using 2 (object: low vs. high) by 2 (probe: low vs. high) univariate, repeated-measures ANOVAs (α = .05). In the case of a significant interaction, one-tailed paired-t-tests Bonferroni-corrected for multiple comparisons (α = 0.0125) were performed separately on the two probe stimulation frequencies. Possible effects of object texture (low vs. high texture frequency) and probe stimulation (low vs. high stimulation frequency) on the kinematic parameters, the force variables and the stimulation time were examined using univariate repeated-measures ANOVAs (α = .05). Effect sizes are reported as Cohen’s d for t-tests and as generalized Eta squared for ANOVAs. The statistical analyses were carried out with R (R Foundation for Statistical Computing, Vienna, Austria).

## Acknowledgments

We thank Aaron Zöller for his advice regarding the software, as well as Daniel Famularo and Leah Trawnitschek for their valuable assistance in data collection. This work was supported by the German Research Foundation (DFG)—Collaborative Research Centre SFB/TRR 135, Project A4 and A5, Grant Number 222641018.

## Author Contributions

Conceptualization, D.V., K.D., K.F.; Methodology, E.F., D.V., A.L., K.D.; Software, E.F., D.V., A.L., K.D.; Investigation, E.F.; Resources, K.F.; Writing –Original Draft, E.F.; Writing –Review & Editing, E.F., D.V., K.F.; Visualization, E.F.; Funding Acquisition, K.F.; Supervision, D.V., K.D., K.F.

## References

1. Voss, M., Ingram, J. N., Haggard, P., & Wolpert, D. M. (2006). Sensorimotor attenuation by central motor command signals in the absence of movement. Nature Neuroscience, 9(1), 26–27. https://doi.org/10.1038/nn1592

2. Chapman, C. E., & Beauchamp, E. (2006). Differential controls over tactile detection in humans by motor commands and peripheral reafference. Journal of Neurophysiology, 96(3), 1664–1675. https://doi.org/10.1139/y94-080

3. Colino, F. L., Buckingham, G., Cheng, D. T., van Donkelaar, P., & Binsted, G. (2014). Tactile gating in a reaching and grasping task. Physiological Reports, 2(3), e00267. https://doi.org/10.1002/phy2.267

4. Wolpert, D. M., & Flanagan, J. R. (2001). Motor prediction. Current Biology, 11(18), R729–R732. https://doi.org/10.1016/S0960-9822(01)00432-8

5. Press, C., Kok, P., & Yon, D. (2020). The perceptual prediction paradox. Trends in Cognitive Sciences, 24(1), 13–24. https://doi.org/10.1016/j.tics.2019.11.003

6. Voudouris, D., & Fiehler, K. (2021). Dynamic temporal modulation of somatosensory processing during reaching. Scientific Reports, 11(1), 1928. https://doi.org/10.1038/s41598-021-81156-0

7. Kilteni, K., & Ehrsson, H. H. (2020). Predictive attenuation of touch and tactile gating are distinct perceptual phenomena. BioRxiv. https://doi.org/10.1101/2020.11.13.381202

8. Seki, K., & Fetz, E. E. (2012). Gating of sensory input at spinal and cortical levels during preparation and execution of voluntary movement. Journal of Neuroscience, 32(3), 890–902. https://doi.org/10.1523/JNEUROSCI.4958-11.2012

9. Bolanowski, S. J., Gescheider, G. A., Verrillo, R. T., & Chechosky, C. M. (1988). Four channels mediate the mechanical aspects of touch. The Journal of the Acoustical Society of America, 84(5), 1680–1694. https://doi.org/10.1121/1.397184

10. Johnson, K. O., Yoshioka, T., & Vega–Bermudez, F. (2000). Tactile functions of mechanoreceptive afferents innervating the hand. Journal of Clinical Neurophysiology, 17(6), 539–558. https://doi.org/10.1097/00004691-200011000-00002

11. Brown, H., Adams, R. A., Parees, I., Edwards, M., & Friston, K. J. (2013). Active inference, sensory attenuation and illusions. Cognitive Processing, 14(4), 411– 427. https://doi.org/10.1007/s10339-013-0571-3

12. Voudouris, D., Broda, M. D., & Fiehler, K. (2019). Anticipatory grasping control modulates somatosensory perception. Journal of Vision, 19(5), 4. https://doi.org/10.1167/19.5.4

13. Voudouris, D., & Fiehler, K. (2017). Enhancement and suppression of tactile signals during reaching. Journal of Experimental Psychology: Human Perception and Performance, 43(6), 1238–1248. https://doi.org/10.1037/xhp0000373

14. Baguley, T. (2012). Calculating and graphing within-subject confidence intervals for ANOVA. Behavior Research Methods, 44(1), 158–175. https://doi.org/10.3758/s13428-011-0123-7

15. Morey, R. D. (2008). Confidence intervals from normalized data: A correction to Cousineau (2005). Tutorial in Quantitative Methods for Psychology, 4(2), 61–64.

16. Fraser, L. E., & Fiehler, K. (2018). Predicted reach consequences drive time course of tactile suppression. Behavioural Brain Research, 350, 54–64. https://doi.org/10.1016/J.BBR.2018.05.010

17. Chapman, C. E., Bushnell, M. C., Miron, D., Duncan, G. H., & Lund, J. P. (1987). Sensory perception during movement in man. Experimental Brain Research, 68(3), 516–524. https://doi.org/10.1007/BF00249795

18. Voss, M., Ingram, J. N., Wolpert, D. M., & Haggard, P. (2008). Mere Expectation to Move Causes Attenuation of Sensory Signals. PLoS ONE, 3(8), e2866. https://doi.org/10.1371/journal.pone.0002866

19. Colino, F. L., & Binsted, G. (2016). Time course of tactile gating in a reach-to-grasp and lift task. Journal of Motor Behavior, 48(5), 390–400. https://doi.org/10.1080/00222895.2015.1113917

20. Bäß, P., Jacobsen, T., & Schröger, E. (2008). Suppression of the auditory N1 event-related potential component with unpredictable self-initiated tones: Evidence for internal forward models with dynamic stimulation. International Journal of Psychophysiology, 70(2), 137–143. https://doi.org/10.1016/J.IJPSYCHO.2008.06.005

21. Martikainen, M. H., Kaneko, K., & Hari, R. (2004). Suppressed responses to self-triggered sounds in the human auditory cortex. Cerebral Cortex, 15(3), 299–302. https://doi.org/10.1093/cercor/bhh131

22. Heinks-Maldonado, T. H., Mathalon, D. H., Gray, M., & Ford, J. M. (2005). Fine-tuning of auditory cortex during speech production. Psychophysiology, 42(2), 180– 190. https://doi.org/10.1111/j.1469-8986.2005.00272.x

23. Cardoso-Leite, P., Mamassian, P., Schütz-Bosbach, S., & Waszak, F. (2010). A new look at sensory attenuation: Action-effect anticipation affects sensitivity, not response bias. Psychological Science, 21(12), 1740–1745. https://doi.org/10.1177/0956797610389187

24. Gremmler, S., & Lappe, M. (2017). Saccadic suppression during voluntary versus reactive saccades. Journal of Vision, 17(8), 8. https://doi.org/10.1167/17.8.8

25. Palmer, C. E., Davare, M., & Kilner, J. M. (2016). Physiological and perceptual sensory attenuation have different underlying neurophysiological correlates. Journal of Neuroscience, 36(42), 10803–10812. https://doi.org/10.1523/JNEUROSCI.1694-16.2016

26. Franklin, D. W., & Wolpert, D. M. (2011). Computational mechanisms of sensorimotor control. Neuron, 72, 425–442. https://doi.org/10.1016/j.neuron.2011.10.006

27. Franklin, S., Wolpert, D. M., & Franklin, D. W. (2012). Visuomotor feedback gains upregulate during the learning of novel dynamics. Journal of Neurophysiology, 108(2), 467–478. https://doi.org/10.1152/jn.01123.2011

28. Körding, K. P., & Wolpert, D. M. (2004). Bayesian integration in sensorimotor learning. Nature, 427(6971), 244–247. https://doi.org/10.1038/nature02169

29. Arikan, B. E., Voudouris, D., Voudouri-Gertz, H., Sommer, J., & Fiehler, K. (2021). Reach-relevant somatosensory signals modulate activity in the tactile suppression network. NeuroImage, 236, 118000. https://doi.org/10.1016/j.neuroimage.2021.118000

30. Manzone, D. M., Inglis, J. T., Franks, I. M., & Chua, R. (2018). Relevance-dependent modulation of tactile suppression during active, passive and pantomime reach-to-grasp movements. Behavioural Brain Research, 339, 93–105. https://doi.org/10.1016/j.bbr.2017.11.024

31. Gertz, H., Voudouris, D., & Fiehler, K. (2017). Reach-relevant somatosensory signals modulate tactile suppression. Journal of Neurophysiology, 117(6), 2262–2268. https://doi.org/10.1152/jn.00052.2017

32. Cybulska-Klosowicz, A., Meftah, E. M., Raby, M., Lemieux, M. L., & Chapman, C. E. (2011). A critical speed for gating of tactile detection during voluntary movement. Experimental Brain Research, 210(2), 291–301. https://doi.org/10.1007/s00221-011-2632-0

33. Broda, M. D., Fiehler, K., & Voudouris, D. (2020). The influence of afferent input on somatosensory suppression during grasping. Scientific Reports, 10(1), 18692. https://doi.org/10.1038/s41598-020-75610-8

34. Dimitriou, M. (2014). Human Muscle Spindle Sensitivity Reflects the Balance of Activity between Antagonistic Muscles. The Journal of Neuroscience, 34(41), 13644–13655. https://doi.org/10.1523/JNEUROSCI.2611-14.2014

35. Friston, K. (2012). A free energy principle for biological systems. Entropy, 14, 2100–2121. https://doi.org/10.3390/e14112100

36. Friston, K. (2009). The free-energy principle: a rough guide to the brain? Trends in Cognitive Sciences, 13(7), 293–301. https://doi.org/10.1016/j.tics.2009.04.005

37. Yon, D., & Press, C. (2017). Predicted action consequences are perceptually facilitated before cancellation. Journal of Experimental Psychology: Human Perception and Performance, 43(6), 1073–1083. https://doi.org/10.1037/xhp0000385

38. Bays, P. M., Flanagan, J. R., & Wolpert, D. M. (2006). Attenuation of Self-Generated Tactile Sensations Is Predictive, not Postdictive. PLoS Biology, 4(2), e28. https://doi.org/10.1371/journal.pbio.0040028

39. Williams, S. R., & Chapman, C. E. (2002). Time course and magnitude of movement-related gating of tactile detection in humans. III. Effect of motor tasks. Journal of Neurophysiology, 88(2), 1968–1979. https://doi.org/10.1152/jn.00527.2001

40. Humes, L. E., Busey, T. A., Craig, J. C., & Kewley-Port, D. (2009). The effects of age on sensory thresholds and temporal gap detection in hearing, vision, and touch. Attention, Perception, & Psychophysics, 71(4), 860–871. https://doi.org/10.3758/APP

41. Diamond, M. R., Ross, J., & Morrone, M. C. (2000). Extraretinal control of saccadic cuppression. The Journal of Neuroscience, 20(9), 3449–3455.

42. Schröger, E., Marzecová, A., & Sanmiguel, I. (2015). Attention and prediction in human audition: A lesson from cognitive psychophysiology. European Journal of Neuroscience, 41(5), 641–664. https://doi.org/10.1111/ejn.12816

43. Railo, H., Olkoniemi, H., Eeronheimo, E., Pääkkönen, O., Joutsa, J., & Kaasinen, V. (2018). Dopamine and eye movement control in Parkinson’s disease: deficits in corollary discharge signals? PeerJ, 6, e6038. https://doi.org/10.7717/peerj.6038

44. Belayachi, S., & Van der Linden, M. (2010). Feeling of doing in obsessive-compulsive checking. Consciousness and Cognition, 19(2), 534–546. https://doi.org/10.1016/j.concog.2010.02.001

45. Lindner, A., Thier, P., Kircher, T. T. J., Haarmeier, T., & Leube, D. T. (2005). Disorders of agency in schizophrenia correlate with an inability to compensate for the sensory consequences of actions. Current Biology, 15(12), 1119–1124. https://doi.org/10.1016/j.cub.2005.05.049

46. Ford, J. M., Palzes, V. A., Roach, B. J., & Mathalon, D. H. (2014). Did I do that? Abnormal predictive processes in schizophrenia when button pressing to deliver a tone. Schizophrenia Bulletin, 40(4), 804–812. https://doi.org/10.1093/schbul/sbt072

47. Yaple, Z. A., Tolomeo, S., & Yu, R. (2021). Abnormal prediction error processing in schizophrenia and depression. Human Brain Mapping, 42, 3547–3560. https://doi.org/10.1002/hbm.25453

48. Oldfield, R. C. (1971). The assessment and analysis of handedness: The Edinburgh Inventory. Neuropsychologia, 9, 97–113. https://doi.org/10.1016/0028-3932(71)90067-4

49. Lakens, D., & Caldwell, A. R. (2021). Simulation-based power analysis for factorial Analysis of Variance designs. Advances in Methods and Practices in Psychological Science, 4(1). https://doi.org/10.1177/2515245920951503

50. Schot, W. D., Brenner, E., & Smeets, J. B. J. (2010). Robust movement segmentation by combining multiple sources of information. Journal of Neuroscience Methods, 187(2), 147–155. https://doi.org/10.1016/j.jneumeth.2010.01.004

51. Laskin, S. E., & Spencer, W. A. (1979). Cutaneous masking. II. Geometry of excitatory andinhibitory receptive fields of single units in somatosensory cortex of the cat. Journal of Neurophysiology, 42(4), 1061–1082. https://doi.org/10.1152/jn.1979.42.4.1061

52. Schütt, H. H., Harmeling, S., Macke, J. H., & Wichmann, F. A. (2016). Painfree and accurate Bayesian estimation of psychometric functions for (potentially) overdispersed data. Vision Research, 122, 105–123. https://doi.org/10.1016/j.visres.2016.02.002

