## Supplement for "Tactile suppression stems from sensation-specific sensorimotor predictions"

**Table**

*Peak-to-peak displacements of the probe stimuli*

| 40 Hz | 140 Hz | 240 Hz |
|---|---|---|
| 7.18 µm | 15.17 µm | 1.61 µm |
| 15.46 µm | 32.95 µm | 3.23 µm |
| 23.22 µm | 50.82 µm | 4.84 µm |
| 30.99 µm | 67.50 µm | 6.46 µm |
| 38.79 µm | 88.14 µm | 8.07 µm |
| 46.61 µm | 106.30 µm | 9.69 µm |
| 54.45 µm | 123.25 µm | 11.31 µm |
